# FlowDesign: Improved Design of Antibody CDRs Through Flow Matching and Better Prior Distributions

**DOI:** 10.1101/2024.11.07.622422

**Authors:** Jun Wu, Xiangzhe Kong, Ningguan Sun, Jing Wei, Sisi Shan, Fuli Feng, Feng Wu, Jian Peng, Linqi Zhang, Yang Liu, Jianzhu Ma

**Affiliations:** University of Science and Technology of China, Hefei, China; Department of Computer Science and Technology, Tsinghua University, Beijing, China; Institute for AI Industry Research, Tsinghua University, Beijing, China; Helixon US Inc, USA; Comprehensive AIDS Research Center, Center for Infection Biology, Pandemic Research Alliance, School of Basic Medical Sciences, Tsinghua Medicine, Tsinghua University, Beijing, China; Department of Electronic Engineering, Tsinghua University, Beijing, China

## Abstract

Designing antibodies with desired binding specificity and affinity is essential for pharmaceutical research. While diffusion-based models have advanced the co-design of the Complementarity-Determining Regions (CDRs) sequences and structures, challenges remain, including non-informative prior distribution, incompatibility with discrete amino acid types, and impractical computational cost in large-scale sampling. To address these, we proposed FlowDesign, a sequence-structure co-design approach based on Flow Matching, offering: (1) Flexible selection of prior distributions; (2) Direct matching of discrete distributions; (3) Enhanced computational efficiency for large-scale sampling. By leveraging various priors, data-driven structural models proved the most informative. FlowDesign outperformed baselines in Amino Acid Recovery (AAR), RMSD, and Rosetta energy. We also applied FlowDesign to design antibodies targeting the HIV-1 receptor CD4. FlowDesign yielded antibodies with improved binding affinity and neutralizing potency compared to the antibody Ibalizumab across multiple HIV mutants, validated by Biolayer Interferometry (BLI) and pseudovirus neutralization. This highlights FlowDesign’s potential in antibody and protein design. A record of this paper’s Transparent Peer Review process is included in the Supplemental Information.

## Introduction

Antibodies are essential to the immune system for their ability to neutralize pathogens^1^ by binding to specific antigens. Designing antibodies that can target particular antigens has significant implications for pharmaceutical and biological research^2^. The Complementarity-Determining Regions (CDRs) are pivotal in the functional repertoire of antibodies, as they constitute the majority of the binding area. Therefore, the primary focus of antibody design is to identify the CDRs with favorable binding affinity and specificity to the antigen. However, it is not feasible to traverse the entire combinatorial space of CDRs via laboratory experiments due to the unaffordable cost, thus efficient exploration with computational methodologies becomes crucial.

Conventional methods sample sequences and structures through simulations with empirical energy functions^3–6^, which are time-consuming and prone to be stuck in local optima. Recent progress exploits deep-learning models to design CDRs or the entire antibody structure^7–12^. Protein language models (PLM), akin to their counterparts in natural language processing, offer a transformative methodology for designing antibody sequences^13, 14^. PLM learns the grammar of protein sequences by training on vast datasets of known proteins, thereby capturing the intricate patterns among amino acids that determine protein structure and function. When applied to the task of antibody design, PLM can predict sequences for the CDRs that are likely to bind with high specificity and affinity to target antigens^15–17^. The use of PLM in antibody design accelerates the discovery process, reducing the reliance on exhaustive wet lab experimentation. By efficiently narrowing down the vast search space to a manageable subset of promising candidates, these models facilitate a more focused and cost-effective empirical validation process.

While they showed potential when adapted to the antibody domain^15–17^, the lack of structural modeling made them suboptimal^7, 9^. A critical issue of sequence-based methods is that they can only enhance the stability of antibodies based on the sequence of individual antibodies. They struggle in improving the affinity between antibodies and antigens because they are unable to describe the sequence or structure of antigens. Another category of promising methods is diffusion models for sequence-structure co-design^11^, which not only model the intricate relations between sequences and structures but also enable diverse sampling of antibodies under specified conditions. Diffusion-based models first inject noises into the data in the forward process, resulting in a prior distribution which are usually standard Gaussian, then learn a generative reverse process to reconstruct the data distribution by denoising the noisy states. In the context of antibody design, the application of diffusion models on both sequences and structures of the CDRs excels in capturing the joint distribution and generates antibodies with enhanced binding affinity^11, 12, 18^.

Despite the encouraging progress, current diffusion-based antibody-design models still suffer from multiple drawbacks. First, the prior distribution, which is commonly a standard Gaussian, fails to reflect the physical plausibility and feasibility of antibody molecules. CDRs consist of continuous linear loop structures constrained by structurally conserved frameworks, with certain position-specific amino-acid preferences. While other deep-learning-based^8, 15, 17^ or energy-based models^3–6^ can easily integrate these prior knowledge into their model initialization, current diffusion models overlook these aspects with the prior distribution limited to standard Gaussian distribution. Second, the continuous nature of the diffusion algorithm is barely compatible with the discrete nature of amino acid types. Adding noises leads to smooth transitive processes but applying it to amino acid types easily induces abrupt changes between consecutive steps, which hinders the model from capturing the desired sequence distribution precisely. Third, the iterative denoising procedure incurs impractical computational costs for large-scale sampling. Diffusion models generate each sample through *T* denoising steps, necessitating running the model for multiple iterations. Therefore, generating large-scale samples for screening potential candidates becomes computationally challenging due to the increased computational cost.

To address the issues, we developed a algorithm, named FlowDesign, to tackle antibody sequence-structure co-design tasks based on Flow Matching^19^. Unlike traditional diffusion-based models, our model approached CDR design as a Transport Mapping problem, which learned a direct mapping from arbitrary initial distributions to the target distribution. Resembling a one-step “diffusion” model, it naturally enjoys the following desirable merits: (1) Flexibility in the selection of the prior distribution, which allows integration of diverse prior knowledge; (2) Direct matching of discrete distributions, which avoids non-smooth generative processes on the amino acid types; (3) High efficiency, which facilitates large-scale sampling with low computational cost. We first assessed the impact of prior knowledge in different forms, including distributions derived from protein language models, physicochemical functions, and structural data-driven models^7^, among which the structural data-driven models yielded the best prior distribution. Next, we conducted a comprehensive evaluation of the sequences and structures of antibodies designed by different computational methodologies. Results demonstrated that adding proper prior knowledge could almost always improve performance with respect to different metrics. In addition, we evaluated the performance of FlowDesign in designing antibodies at both the molecular and cellular levels against Human Immunodeficiency Virus Type 1 (HIV-1). HIV-1 has caused a significant threat to human health. Broadly neutralizing antibodies (bnAbs) against HIV-1 are promising candidates for HIV-1 prevention and treatment. However, high-frequency mutations and a large number of resistant strains of HIV-1 make it hard to discover antibodies through traditional antibody screening methods. Therefore, there’s an urgent need to use the AI model to assist in the design of antibodies against HIV-1. We used the FlowDesign model to redesign the HCDR3 of the HIV-1 antibody Ibalizumab (imab), which targets the HIV-1 cellular receptor CD4. We evaluated the designed antibodies through SPR and HIV-1 pseudovirus neutralization assay and found multiple candidates have comparable or even better binding affinity and neutralizing compared with the state-of-the-art HIV antibody Ibalizumab^20^. These results showed that our model has the ability to regenerate antibodies and has potential applications in engineering other protein molecules.

## Results

### Overview of FlowDesign

We represented an antibody as {**s**_*i*_, **x**_*i*_, **O**_*i*_}, namely the amino acid type, the *C*_*α*_ atom coordinate and the backbone orientation of each residue in the antibody(Figure 1.A). The inference process begins with a given antibody-antigen complex with CDRs to be designed. The CDRs are initialized from a certain prior distribution. There are different options for prior distributions, which we discuss in the next section. Subsequently, the model predicts drift forces on each residue from the inputs, which are interpreted as the transformation from the initial distribution to the target distribution. Given an informative initial distribution, we can directly perform a one-step transformation to achieve the target distribution. However, if delicate refinements are required, we can also simulate an integration process by scaling the predicted drift force with a coefficient and repeating the cycle for *N* times, where the selection of *N* depends on the application scenarios. After the inference, we adopted the side-chain packing function in the Rosetta package^5^ to construct the side chains and perform relaxation to prevent inter-atomic collisions. As the side-chain packing software we adopted only produces a small portion of atom clashes. During the sampling stage, we directly discarded structures with atomic clashes. As for training, we exerted supervision on the drift forces to update the parameters of the model. Specifically, for each antibody-antigen complex in the dataset, we calculated its difference from an initial complex sampled from the prior distribution as the ground truth for drift forces. Then we minimized the distance between the predicted drift force and the ground truth one given the same initial complex. Unlike other diffusion-based models, which repeatedly add and remove noise, our training objective is a one-step transformation, directly matching the initial distribution with the target distribution. FlowDesign offers two design approaches: The first mode designs the antibody CDR given the framework and antigen complex. The second mode fixes the framework’s amino acid type and structure, employs dyMEAN^8^ and AlphaFold3^21^ for complex prediction, and subsequently designs the CDRs.

**Figure 1.**
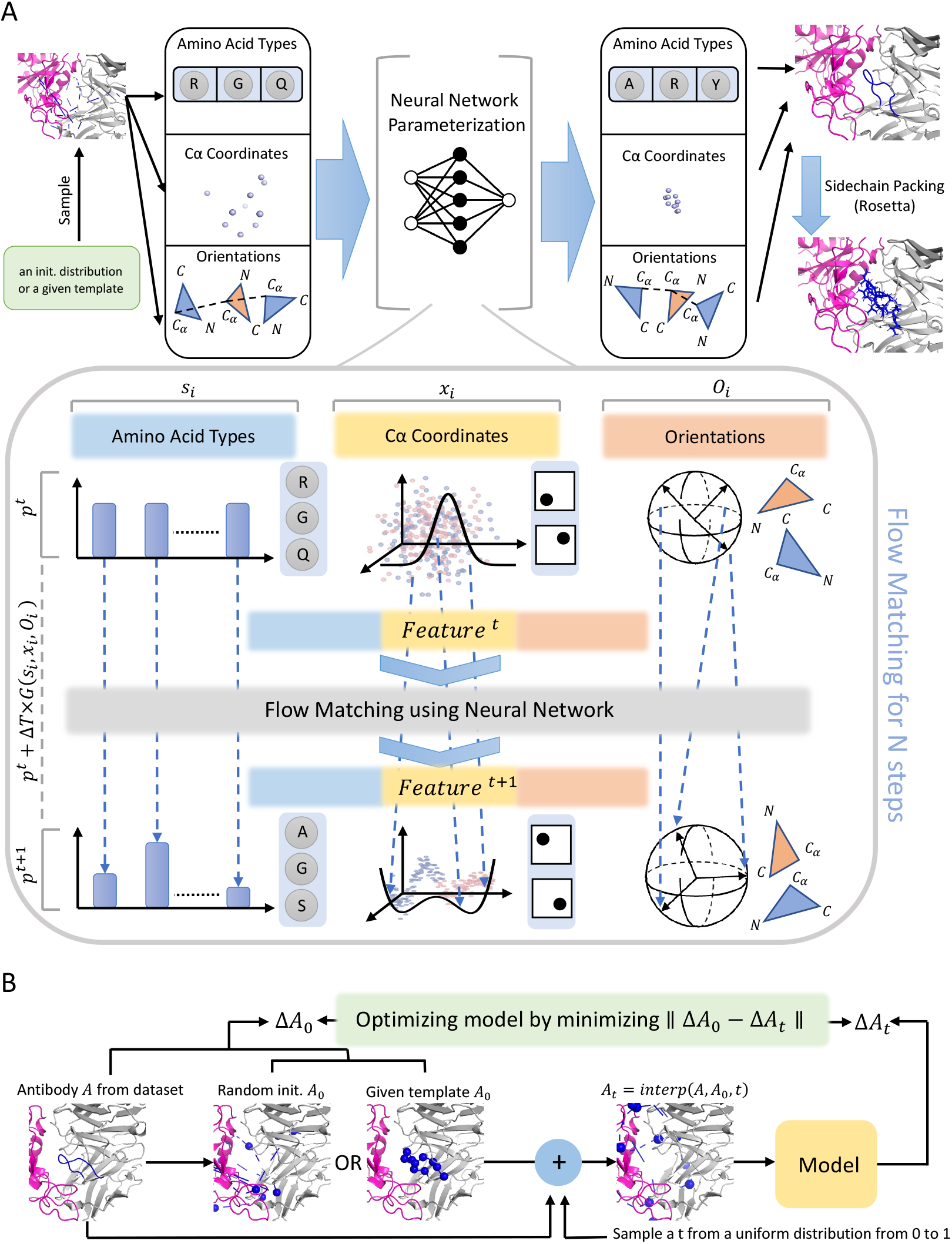
Inference and training process of the FlowDesign model. **(A)** FlowDesign learns to generate antibodies by matching the prior distribution to the data distribution extracted from the antibody dataset. **(B)** In the training process, the model parameters were trained by minimizing the difference between the initialized distribution and the target distribution.

### Performance for different prior distributions

In contrast to conventional diffusion-based models, our model enjoys the flexibility to integrate various types of prior distribution into the generation process. To assess the impacts of different prior distributions and find the optimal strategy, we conducted attempts on four distinct prior distributions: random distribution, the sequence-structure joint distribution outputted by an all-atom model dyMEAN^8^, the structure distribution generated by the kinematic-closure-based method KIC^22, 23^, and the sequence distribution outputted by a large protein language model ESM2^13^. The model was trained and validated on CDRH3 with the four prior distributions because of the critical role in antigen-antibody binding and the high diversity of CDRH3. Experimental results revealed that using dyMEAN as the prior distribution generated antibodies with the best quality. In particular, the model utilizing dyMEAN achieved the highest sequence recovery rate, showing a nearly 15% improvement over methods using random initialization (Figure 2A). Protein language model ESM2 struggled with the high diversity of its generated antibody sequences, leading to a lower sequence recovery rate. It could not provide effective prior knowledge for the modeling of antibody structures due to the lack of structural information. The model with dyMEAN prior obtained the lowest average RMSD (root-mean-square deviation), at 2.291Å on average, consistently outperforming the other three methods across various sampling ratios (Figure 2B). This indicated that the predicted conformations were the closest to the original antibody structures. KIC initialization reached relatively poor performance among all the methods, due to the limited number of solutions provided by the kinematic equations due to the unaffordably high computational cost (STAR Methods). In addition, due to the effective sequence and structural knowledge provided by dyMEAN initialization, the generated CDRs had lower energy compared to those from other methods, resulting in a more stable overall binding (Figure 2C). We further provided the visual results of antibody-antigen complexes generated by the dyMEAN-initialized model (Figure 2D), which exhibited better stability compared with results from other methods.

**Figure 2.**
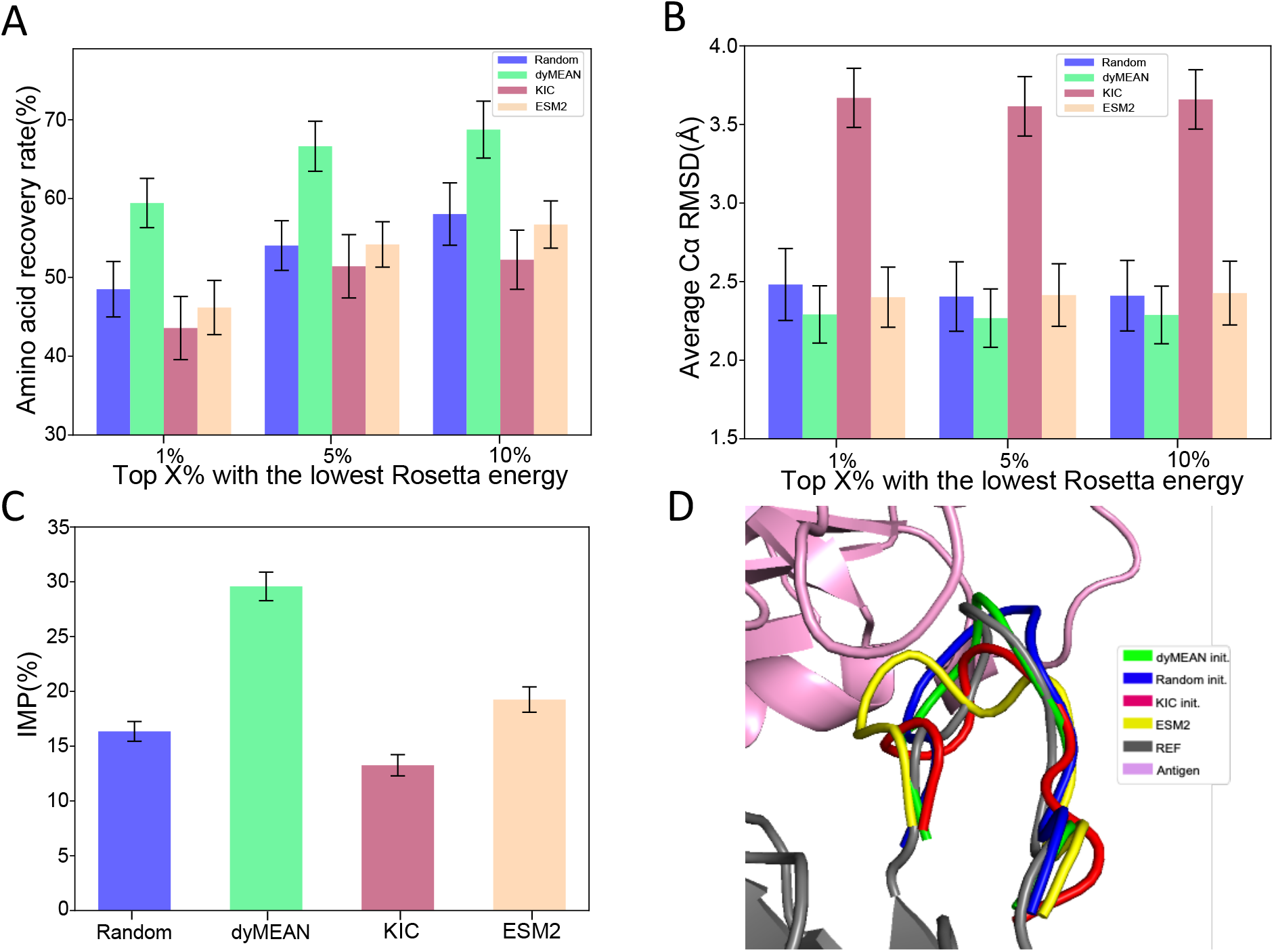
Performance for different prior distributions. **(A)-(C)** The performance of amino acid recovery rate, RMSD, and IMP for antibodies generated by different prior distributions, respectively. IMP represents the proportion of generated antibodies with energy lower than that of the original antibodies. Error bars represent 90% confidence intervals. **(D)** 3D structures of the antibody CDRH3 generated by different initialization methods, where the antigen structure is shown in pink and the original antibody structure is depicted in gray.

### Performance for Sequence-Structure Co-design

To validate the superiority of our model, we compared it against multiple strong baselines for antibody design, including HERN^10^, Refinegnn^9^, RosettaAb^5^ which is based on empirical energies, the diffusion-based model Diffab^11^, and dyMEAN^8^, across diverse metrics. We removed all structural and sequence data for the antibody CDRs and simultaneously designed all CDRs in all H-chains and L-chains together. Notably, the RefineGNN and HERN algorithms do not accept light chains as input, and HERN can only generate sequences. Therefore, we did not test the metrics of these methods on light chains. The experimental results indicated that our model achieved a higher amino acid recovery rate on CDRs compared to other methods due to the effective incorporation of prior knowledge (Figures 3A-F). Notably, our model surpassed other methods by a large margin on CDRH3, which is also the most critical part of binding among all the CDRs. By selecting the top 1% of sampled antibodies based on energy, the amino acid recovery rate on CDRH3 could exceed 60%, which exhibited inspiring potential for enhanced efficacy and efficiency in practical applications. We noticed that our model’s performance improvement is limited at different sampling ratios, with CDRH3 amino acid recovery rate increasing by less than 5% when expanding the sampling range. This is because we achieved sufficiently good conformations within the top 1% of the screening. Expanding the screening range did not improve the metrics, demonstrating that our method can efficiently perform large-scale screening using energy criteria. Moreover, our model achieved a lower average RMSD on CDRs compared to most other methods except dyMEAN. The reason is that the original version of dyMEAN only generates one antibody while FlowDesign is a generative model which can generate multiple antibodies. Given the widespread adoption of high-throughput display platforms, tools that enable the simultaneous design of multiple diverse antibodies are better suited to the practical demands of the biopharmaceutical industry. Specifically, it performed notably better on CDRH3, with an average reduction of 0.693Å compared to Diffab method (Figure 3G). We visualized an example of design by our model in Figure 3H, which illustrated that our model effectively transformed the initial distribution to the target distribution, resulting in an antibody-antigen complex that was more stable than the initial one.

**Figure 3.**
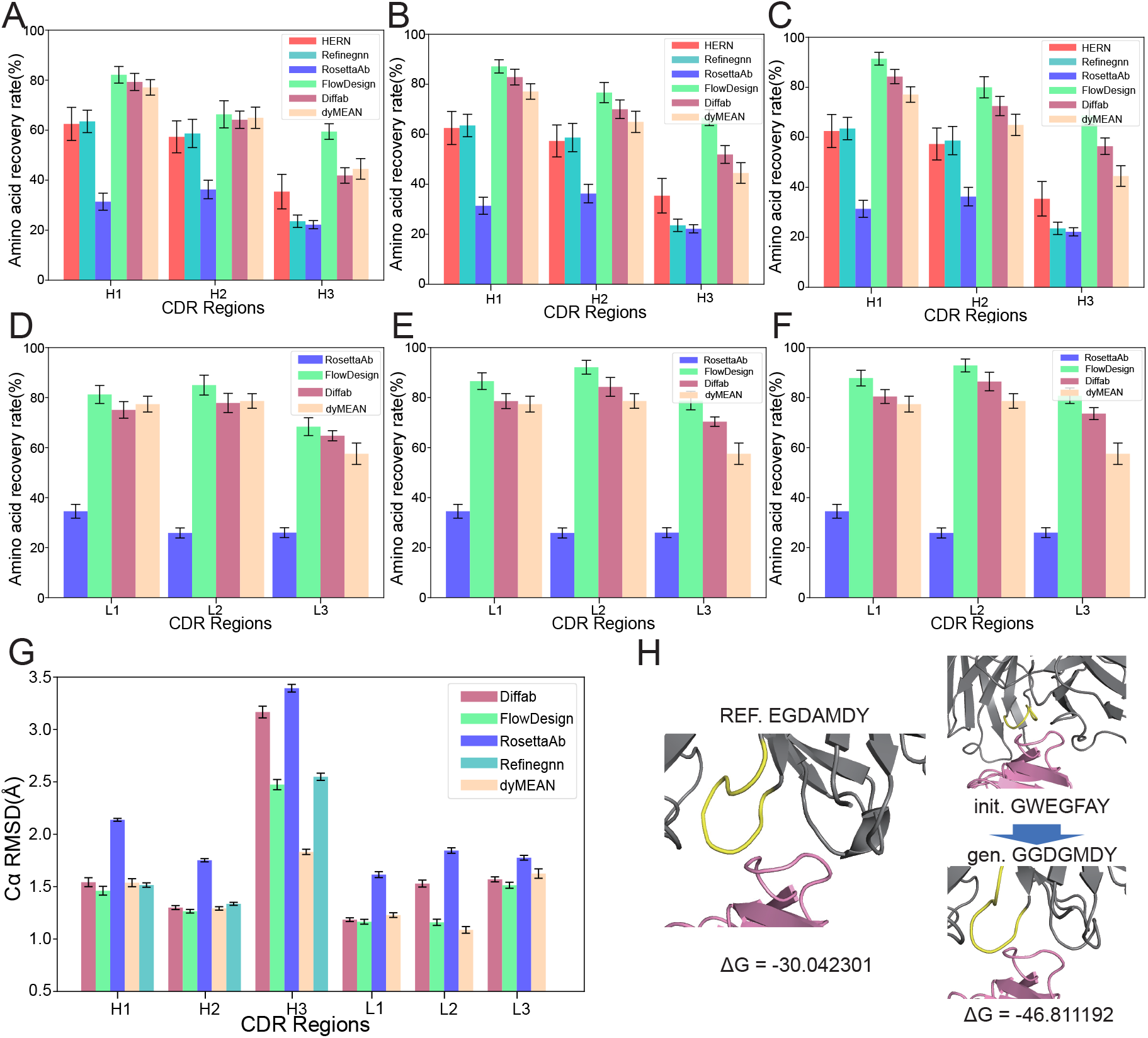
**(A)-(C)** Performance of different models on the amino acid recovery rate in different CDRs in heavy chains. From left to right, antibodies sampled are sorted by the Rosetta energy, and the maximum amino acid recovery rate is calculated for the top 1%, 5%, and 10% of these sampled antibodies. **(D)-(F)** Performance for different CDRs in light chains. From left to right, antibodies sampled are sorted by the Rosetta energy, and the maximum amino acid recovery rate is calculated for the top 1%, 5%, and 10% of these sampled antibodies. **(G)** Average *C*_*α*_ atom RMSD of the antibody structures generated by different models across various CDRs. Error bars represent 90% confidence intervals. **(H)** An example of a prediction made by our model. The left is the original antibody structure, and the right is the antibody predicted by the model from the initial distribution. The gray area represents the antibody framework, the yellow area is the CDRH3 to be generated, and the pink area is the antigen. Δ*G* refers to the binding energy between the antigen and the antibody.

Next, we conducted an extensive assessment of the antibodies generated by our model and Diffab, the only models possessing enough diversity in large-scale sampling among all the models. The antibodies sampled by our model exhibited a lower energy landscape across all CDRs compared to those sampled by Diffab (Figures 4A,B). Additionally, the proportion of antibodies designed with lower energy than the native one is also higher, particularly with a 13.11% increase in CDRH1. This suggested our model was capable of providing more potential candidates than the diffusion-based model when the number of samples was limited. We showed the proportion of designed antibodies with better amino acid recovery than the prior distribution, namely, the distribution from dyMEAN, after energy ranking (Figure 4C). Apart from confirming the superiority of our model over Diffab, the results also confirmed that our model effectively integrated the prior knowledge provided and learned a more complete and accurate distribution based on prior knowledge. In addition, it could be observed that high-quality antibodies were more concentrated at the forefront of the energy ranking. Further, the point-wise comparison suggested that at different energy sampling ratios, our model consistently outperformed the Diffab in the restoration of both amino acids and structures (Figures 4D,E). The experimental results above demonstrated that our model surpassed other models in terms of designing stable and plausible CDRs, which was particularly notable for the typically challenging CDRH3. Furthermore, under the setting of large-scale sampling, our model was capable of yielding more candidates on the desirable energy landscape than Diffab within the same number of samples, exhibiting greater potential for application.

**Figure 4.**
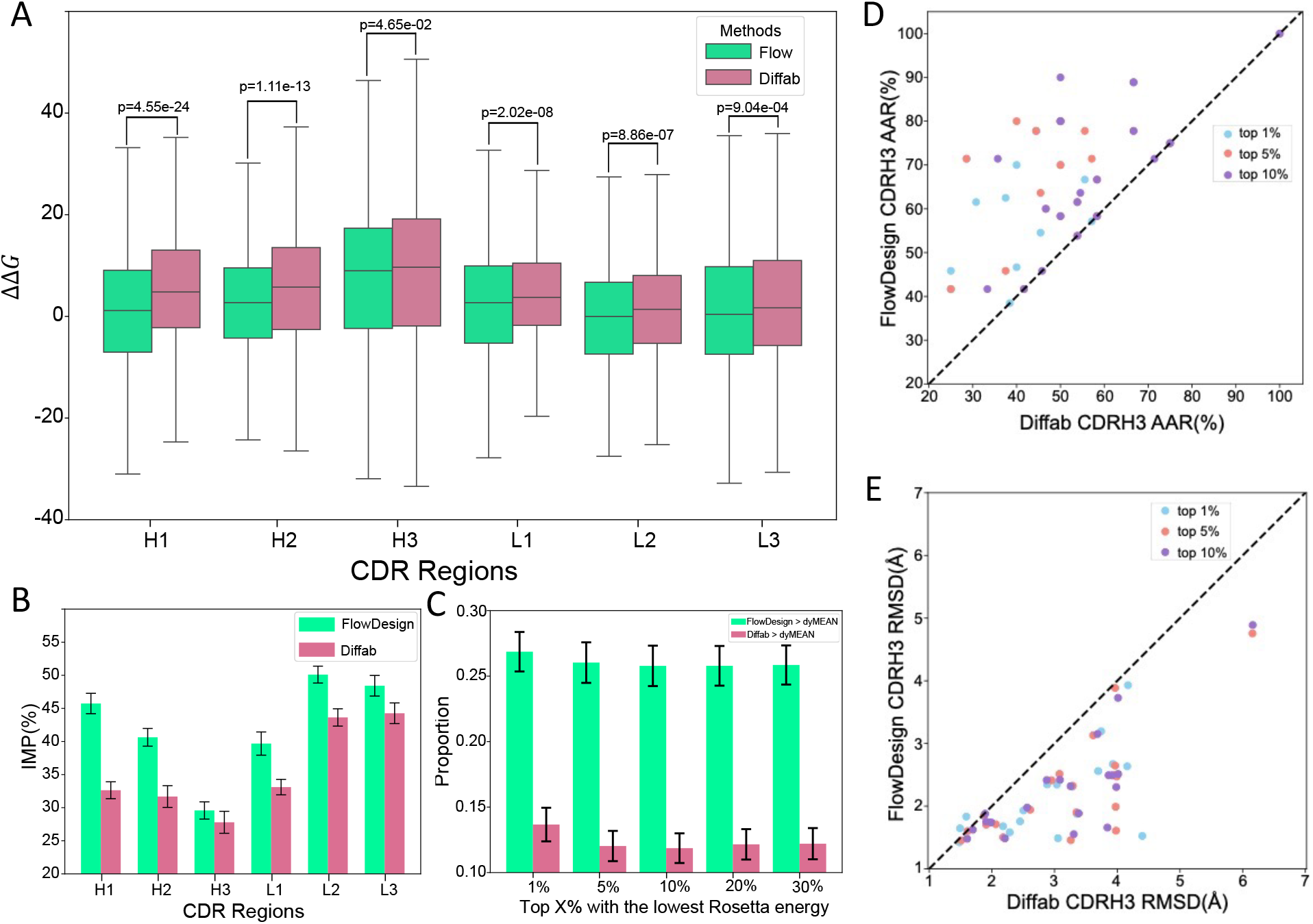
**(A)** The energy (The ΔΔ*G*) difference between the sampled conformation and the original native conformation. **(B)** The proportion of antibodies sampled by the two models that have lower energy compared to the original conformation. **(C)** The proportion of samples from FlowDesign and the diffusion-based model that have the amino acid recovery rate greater than that of the dyMEAN model. **(D)**,**(E)** Performance comparison between FlowDesign and Diffab ranked by Rosetta energy. Error bars represent 90% confidence intervals. P values were computed based on a one-sided paired t-test with n=1000, where n represents the sample size.

We explored the relationship between the RMSD of generated CDRs backbones and the energy of the final conformations. The trending in Figure S3A revealed that higher RMSD of the backbone corresponds to increased energy after side-chain packing. Two generated conformations with high and low backbone RMSD were visualized in Figures S3J,K. It demonstrated that lower backbone RMSD led to lower energy after side-chain packing and better side-chain alignment with the reference data. We also analyzed the influence of backbone RMSD on the side-chain atomic clashes. Results in Figure S3C indicated that the proportion of side-chain clashes during generation was barely related to the type of CDRs. Figures S3D-I visualized the relationship between backbone RMSD and side-chain clashes on different CDRs (i.e. H1, H2. H3, L1, L2, and L3), which did not show significant correlations.

### Performance for designing CDRs based solely on sequences

In real-world scenarios of antibody design and optimization, the specific structure of an antibody might be unknown, and only the sequence is available. To further assess the potential of our method in designing antibody CDRs, we extended our method to design CDRs based solely on the sequence information of the antibody framework. The specific approach involves generating the structure of the antibody framework, excluding the CDRs, from the known sequence. Then, we used FlowDesign for designing the CDRs. We tested two methods to generate the structure: dyMEAN^8^ and AlphaFold3.^21^ We then compared our method, which utilizes known framework structures, with the sequences-based design method across various metrics. Figures 5A, B show the AAR and RMSD of CDRs sampled at different ratios based on energy. It is evident that, aside from cases where the true structure is known, using AlphaFold3 for framework structure prediction framework results in the most accurate sequences and structures for the generated CDRs. Similarly, Figures 5C, G indicate that the energy of structures generated by this method is relatively low, making them well-suited for large-scale sampling. Of most, the most optimal performance is achieved with the known real structure. Among sequences-based design methods, complexes generated using AlphaFold3 are more stable than those from dyMEAN, with performance very close to knowing real structures. Figure 5H also shows that the frameworks generated by AlphaFold3 have lower RMSD, further supporting this point. Figures 5I, J, K visualize the complexes generated by three methods: dyMEAN-generated framework structure, the real framework structure, and the framework structure generated by AlphaFold3. The figures show that dyMEAN’s performance in reconstructing structures from sequences is weak, resulting in docking poses that differ from the real poses, leading to higher energy. In contrast, the structures generated by AlphaFold3 closely resemble the actual structures, yielding results that are more aligned with calculations based on the real framework structures. The experimental results indicate that our method can effectively extend from just using the antibody framework sequence, with performance close to scenarios with known real structures. This demonstrates the robustness of our algorithm.

**Figure 5.**
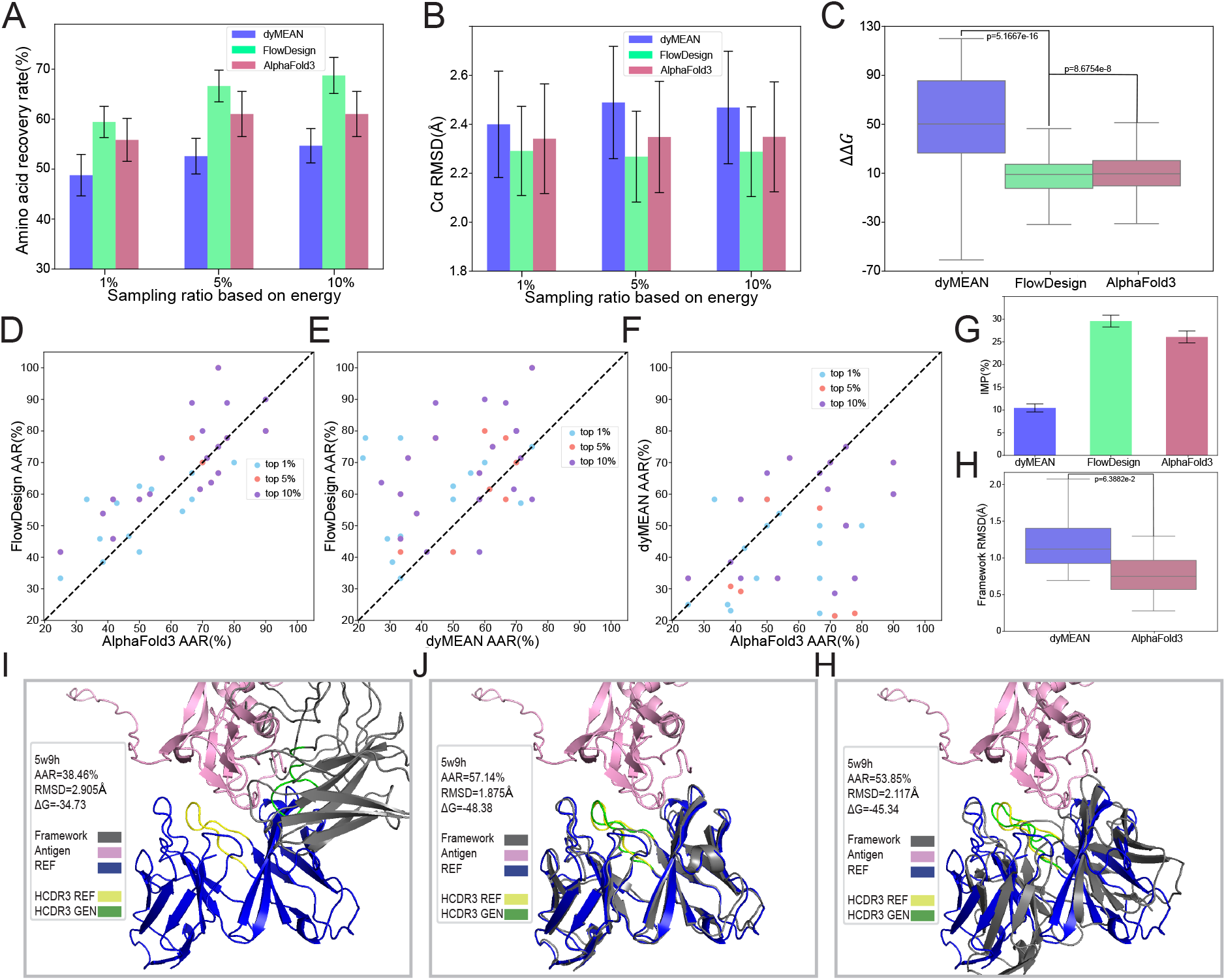
**(A)** Amino acid recovery rate of the dyMEAN generated framework, AlphaFold3 generated framework structure, and the known true structure for the FlowDesign method.**(B)** *C*_*α*_ RMSD of the dyMEAN generated framework, AlphaFold3 generated framework structure, and the known true structure for the FlowDesign method. **(C)** ΔΔ*G* of the dyMEAN generated framework, AlphaFold3 generated framework structure, and the known true structure for the FlowDesign method. **(D)(E)(F)** Performance comparison between three different structure initialization methods ranked by Rosetta energy. **(G)** IMP of three structure initialization methods. **(H)** Comparision of RMSD between frameworks generated by dyMEAN and AlphaFold3. **(I)(J)(K)** Visualization of the generated complex using three structure initialization methods. Error bars represent 90% confidence intervals. P values were computed based on a one-sided paired t-test with n=1000, where n represents the sample size.

### Performance for HIV antibody design

As HIV-1 antibody screening and design is a big challenge using traditional methods, we tried to use FlowDesign to design HIV-1 antibody. Given that CDRH3 primarily affects antibody binding, we fixed the framework of the HIV-1 antibody and redesigned the CDRH3 of the antibody. We used MMseqs2^24^ to filter out antibodies with more than 50% sequence identity in CDRH3 to Ibalizumab (imab) from the dataset and retrained the model. First, we used the model to sample and generate CDRH3 variants under different noise intensities using the framework of antibody Ibalizumab (imab), producing around 50,000 different CDRH3 sequences. We then ranked the generated antibodies using Rosetta energy and selected 12,000 unique sequences for further experiments. A yeast display library was then constructed with these 12000 HCDR3 variants in the Ibalizumab backbone. Subsequently, fluorescence-activated cell sorting (FACS) was performed on the yeast display library based on antibody expression level and binding affinity to CD4. Increasing proportion of the yeast clones bound to CD4 was observed from 0.19% for the first sort (Figure 6A), to 25.9% for the second (Figure 6B) and to 61.7% for the third sorting (Figure 6C). Yeast displayed single clones Ibalizumab and FZD5 antibody 2919 were act as positive control and negative control (Figure 6D, 6E). After the yeast display, we performed next-generation sequencing on all screened antibodies, selected the top 10 antibodies with the highest reads, and chose the three antibodies with the lowest energy for further experiments, named imab-mut-1, imab-mut-2, and imab-mut-3. The equilibrium dissociation constant (KD) with CD4 for these three antibodies were 0.19 nM, 0.56 nM, and 0.90 nM respectively. We assessed the broad-spectrum neutralization capacity by performing pseudoviral neutralization experiments against 11 HIV-1 pseudoviruses. These pseudoviruses are widely distributed, including subtype A, subtype B, subtype C, subtype CRF01, and subtype CRF07, and span from tier 1 to tier 3. Compared with Ibalizumab, the neutralization IC50 values of these three mutants for different pseudoviruses were comparable by orders of magnitude. All three antibodies had a better neutralization ability than the ancestral antibody to some of the pseudoviruses, but overall, imab-mut-2 performed best, followed by imab-mut-1, and imab-mut-3 performed relatively poorly. In the neutralization of 11 pseudoviruses, imab-mut-2 neutralized half of the pseudoviruses (TRO_11, X2278_C2_B6, BJOX002000_03_2, CNE4, CNE14, SF162) better than imab-WT, as shown by the neutralization IC50 value of imab-mut-2 for CNE4 of 0.006 *µ*g/ml, the IC50 value of the imab-WT is 0.024 *µ*g/ml. Furthermore, the neutralization ability of imab-mut-2 to two of these pseudoviruses (398_F1_F6_20, CNE8) was comparable to that of Ibalizumab.

**Figure 6.**
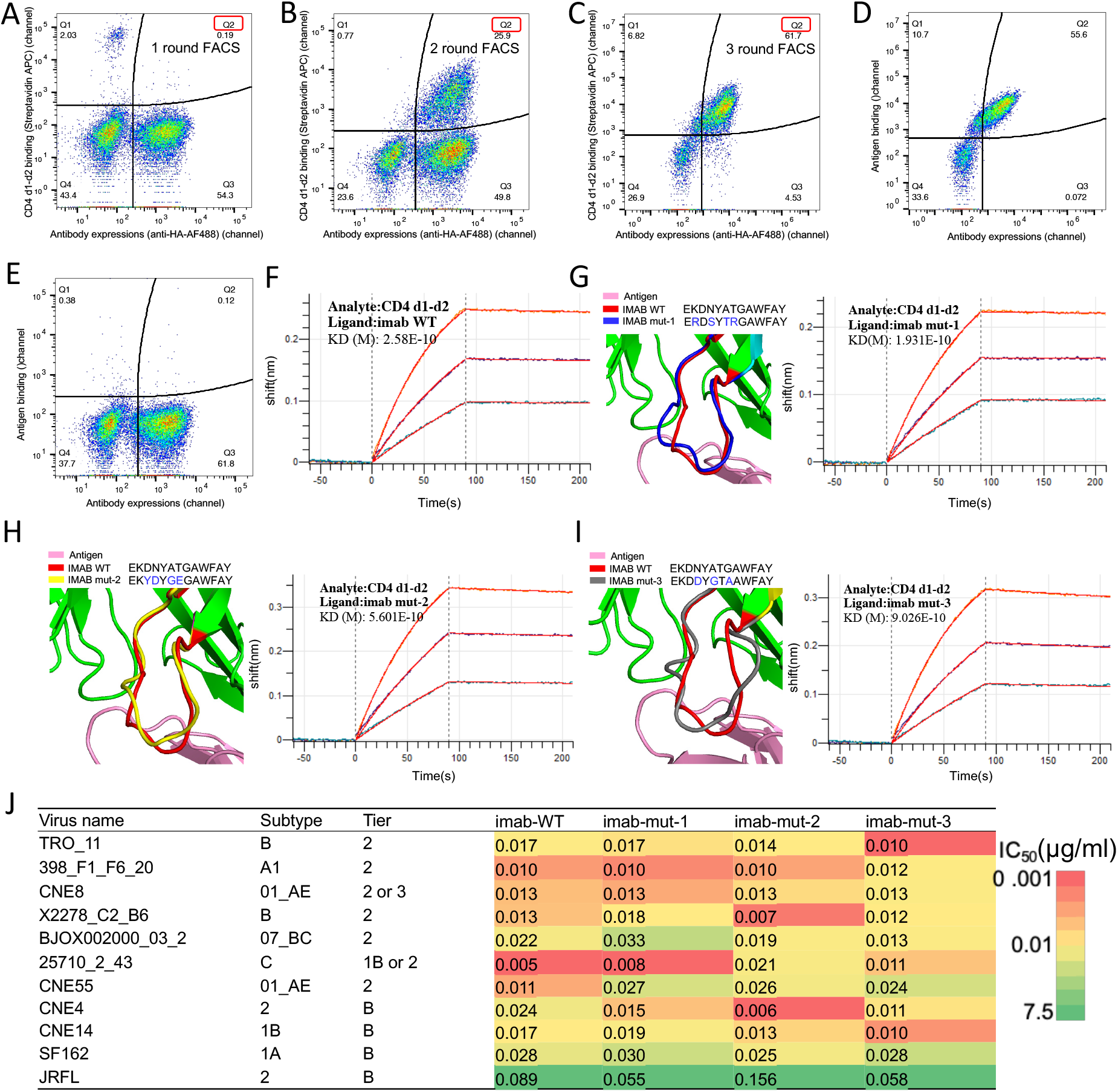
Performance for designed HIV-1 antibody variants. **(A)-(C)** Enrichment progress for designed antibody binding to CD4 through three rounds FACS. Yeast displayed single clones of designed antibodies Ibalizumab and FZD5 antibody 2919 acted as positive control and negative control, respectively. A red box is used to highlight the Q2 area and select its sequence for filtering. **(D, E)** Binding kinetics of Ibalizumab and FZD5. **(F)-(I)** Binding kinetics of Ibalizumab and the top three isolated mAbs imab mut-1, imab mut-2 and imab mut-3 with CD4 d1-d2 measured by Biolayer Interferometry(BLI). The predicted structure of imab-mut1, imab-mut2, and imab-mut3 binding to antigen CD4 also showed in (G)-(I). **(J)** IC50 values of mutated antibodies and imab-WT against pseudotyped HIV-1 strains. Results were calculated from three independent experiments.

**Figure 7.**
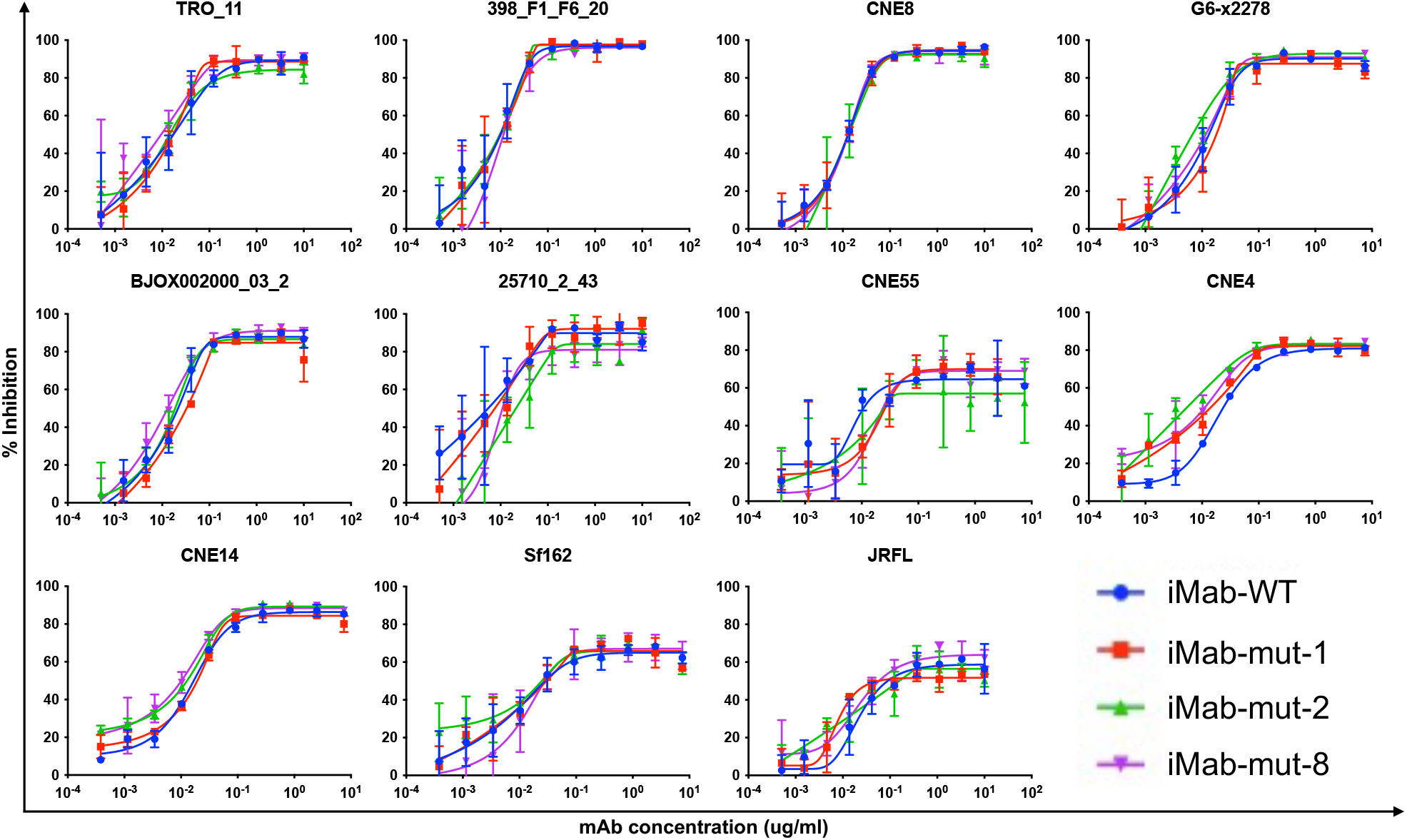
Neutralization curves of the generated antibodies and wild-type Ibalizumab against pseudotyped HIV-1 strains.

We visualized the energy of each residue in the CDRH3 from IMAB-WT and the model samples to identify patterns as shown in Figure S6. It can be observed that the last five amino acids of the CDRH3 (107A, 108W, 109F, 110A, 111Y) region in imab have maintained relatively low energy. Mutations in these five amino acids are likely to increase the conformation’s energy. As shown in Figure S6, the three mutations (mut-1, mut-2, mut-3) with a similar Kd magnitude (1e-10) to imab-WT did not alter this region. However, in Figures S6 D, E, changes at positions 107 and 110 led to a noticeable energy increase in the residues of this region, resulting in an experimentally measured increase in Kd (1e-9). Based on the energy of imab-WT, the residues at positions 103, 104, and 105 show higher energy than their surroundings. This suggests that modifying the residues in this region could potentially improve the binding affinity of imab-WT without affecting the stability of the surrounding regions. Figures S6A-C display three mutations generated by FlowDesign, imab mut-1, mut-2, and mut-3, all of which involve changes in this high-energy segment. It indicates that the energy levels of these residues decrease while the energy of the surrounding regions does not increase significantly. Experimentally, these mutations demonstrate affinities that are on par with or even better than imab-WT.

## Discussion

In this work, we developed a model based on the Flow Matching algorithm which can map any prior distribution to the data distribution to design antibody CDRs. In our algorithm, it is important to note that the model relies on the complex structure of the target antibody and antigen. Incorrect structures may adversely affect the model’s performance. However, recent breakthrough shows that the antibody-antigen complex structures can be accurately predicted given only their protein sequences^21^. Therefore, in the near future, it can be expected that an initial antibody can be obtained either by immunizing animals or using display technology, then the three-dimensional structure of the antibody-antigen complex is predicted by AlphaFold3, and finally the binding affinity could be enhanced through FlowDesign. In our future work, we will focus on how to obtain accurate structures of complexes in specific application scenarios while reducing the model’s dependency on these complexes during inference. We believe this will enhance the applicability of the algorithm in practical antibody design tasks.

## RESOURCE AVAILABILITY

### Lead Contact

Further information and requests for resources and reagents should be directed to and will be fulfilled by the Lead Contact, Jianzhu Ma.

## Materials Availability

This study did not generate new materials.

## Data and Code Availability

- The antibody dataset we used could be downloaded at https://opig.stats.ox.ac.uk/webapps/sabdab-sabpred/sabdab. And its sabdab summary file can be found in our GitHub repository(see key resources table for details).
- All original code has been open-sourced on GitHub. For details, please see the key resources table.
- Any additional information required to reanalyze the data reported in this paper is available from the lead contact upon request.

## Key Resources Table

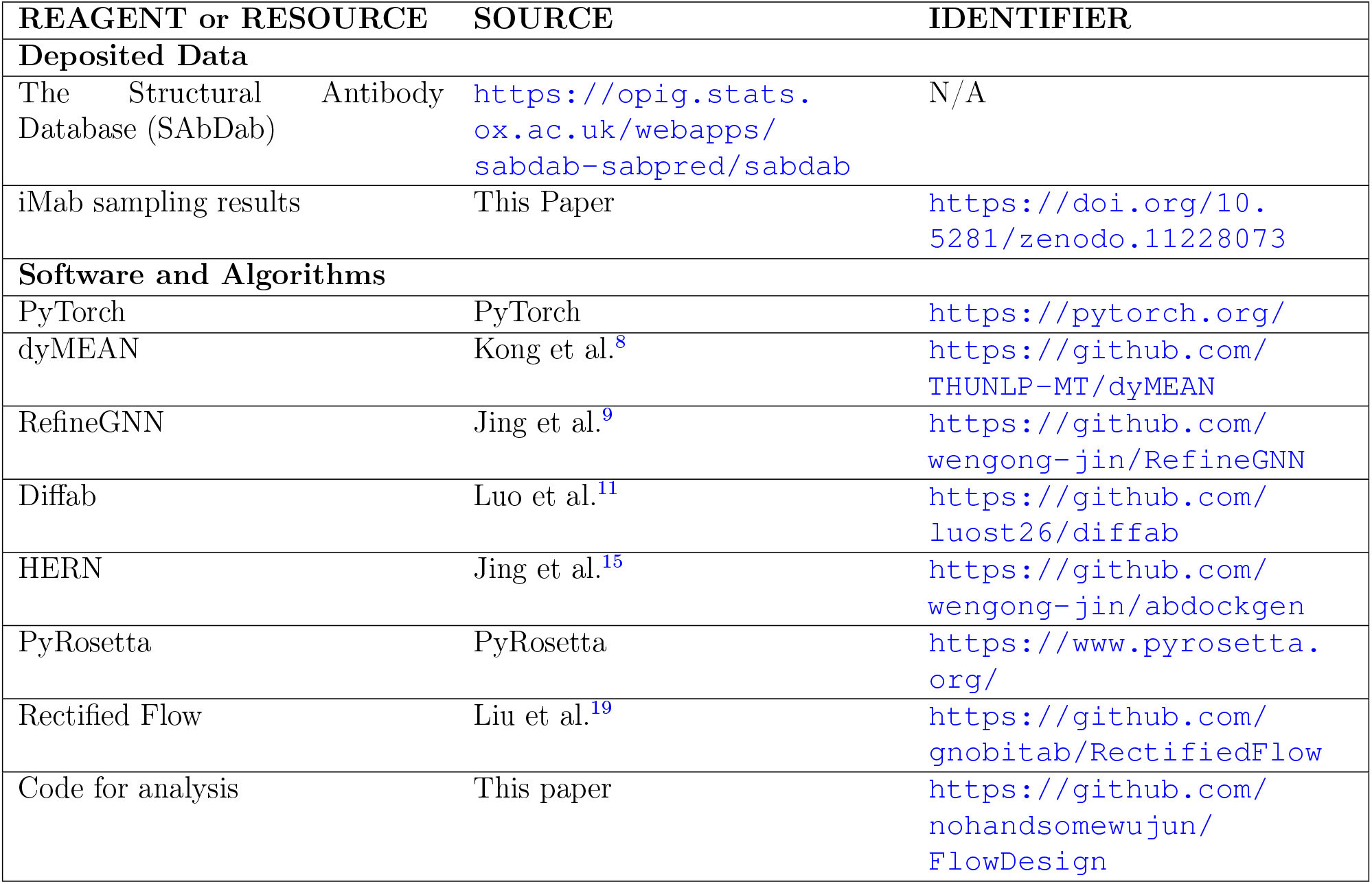

## ACKNOWLEDGMENTS

This work was supported by the National Key Research and Development Program of China grants 2022YFF1203100 and China’s Village Science and Technology City Key Technology funding.

## AUTHOR CONTRIBUTIONS

J.M., Y.L., J.W., X.K., J.P., F.F., and F.W. were involved in the conceptualization of the project. J.W. and X.K. performed the investigation and developed the methodology of the project. N.S., J.W., S.S., and L.Z. conducted the HIV-related biological experiments.

## STAR METHODS

### METHOD DETAILS

In this section, we have explained how to download and run the code for the baselines in Subsection Baselines. We describe the formalization of the antibody CDRs design task from the perspective of the Transport Mapping problem in Subsection Transport Mapping Problem in Antibody CDRs’ design. Next, in Subsection Method of interpolation for antibodies and Subsection Loss Function, we present the method for interpolation of the antibody as well as the loss function for training, respectively. Subsection Parameterization with Neural Network outlines the overall architecture of our model, and Subsection Data Preparation, Subsection Prior Distributions demonstrate the details of the data preparation and the prior distribution for training. Subsection Model Training and Inference provides an overall description of the entire process of model training and inference, along with the selection of hyper-parameters.

### Baselines

When evaluating performance, we selected baselines such as Diffab^11^, RefineGNN^9^, dyMEAN^8^, and HERN^15^. Diffab is a diffusion-based generative model with approximately 4 million parameters, capable of simultaneously sampling and generating a large number of CDR structures and sequences. RefineGNN is an iterative refinement graph neural network for antibody sequence-structure co-design with approximately 6 million parameters. dyMEAN is an end-to-end full-atom model using an adaptive multi-channel equivariant encoder to update both 1D sequences and 3D structures with approximately 2 million parameters. HERN is used to design sequences of CDR via hierarchical equivariant refinement with approximately 7 million parameters. We used the InterfaceAnalyzer-Mover in PyRosetta to calculate binding energy. The weight setting PyRosetta used is REF2015. The binding energy is a weighted sum of various Rosetta energy terms and can be referenced in the paper.^25^ The corresponding code was downloaded from their respective GitHub repositories(see key resources table for details), and we re-partitioned the dataset(see data preparation for details) to retrain these models. In testing generative models like Diffab, we sample the same number of antibodies as the FlowDesign algorithm. For non-generative models, we generate one result for each antibody in the test set.

### Transport Mapping Problem in Antibody CDRs’ design

We begin by presenting an overview of the traditional Transport Mapping Problem:

**Definition 0.1** (Transport Mapping). Given the empirical observations of two distributions *X*_0_ ∼ *π*_0_ and *X*_1_ ∼ *π*_1_ on ℝ ^*d*^, we need to find a transport map *T* : ℝ ^*d*^ → ℝ ^*d*^, such that *Z*_1_ := *T* (*Z*_0_) ∼ *π*_1_ when *Z*_0_ ∼ *π*_0_, that is, (*Z*_0_, *Z*_1_) is a coupling of *π*_0_ and *π*_1_^19^.

The design of antibody CDRs can be similarly conceptualized as a Transport Mapping problem. First, we need to give the computational representation of antibody-antigen complexes as below. Following Diffab^11^, an antibody-antigen complex is represented by the amino acid type *s*_*i*_ ∈ {ACDEFGHIKLMNPQRSTVWY}, the *C*_*α*_ atom coordinate ***x***_***i***_ *∈*ℝ^3^, and the backbone orientation ***O***_***i*** *∈*_ *SO*(3) of each residue, where *i* = 1…*N* for an antibody-antigen complex containing *N* residues. In addition, each *s*_*i*_ is encoded as a one-hot vector ***s***_***i***_ as the input of the model. With the indexes of the residues in the CDR to design from *l* + 1 to *l* + *m*, we denote the CDR as *ℛ* = {***s***_*i*_, ***x***_*i*_, ***O***_*i*_|*i* = *l* + 1, …, *l* + *m*} and the rest of the antibody-antigen complex, which is given as the input, as *C* = {***s***_*j*_, ***x***_*j*_, ***O***_*j*_|*j* ∈ {1, …, *N*}*\*{*l* + 1, …, *l* + *m*}}. The initial state of the CDRs, which is sampled from a specific distribution, is denoted as 𝒯 = {***T*** _***s****i*_, ***T*** _***x****i*_, ***T*** _***O****i*_ |*i* = *l* + 1, …, *l* + *m*}.

Now we are ready to formalize antibody CDR design as a Transport Mapping problem. Given *𝒯*∼ *π*_0|*C*_ and *ℛ*∼ *π*_1|*C*_, we aim to find a transport map *T* such that *T* (*𝒯*)∼ *π*_1|*C*_, where *𝒯*∼ *π* _0|*C*_ and *C* serves as the contextual condition. Obviously, such *T* can help us to generate the target CDR based on the premise of the framework.

### Method of interpolation for antibodies

To endow the model with the capability of fine-grained flow matching, the training process takes the interpolation between the initial distribution and the target distribution as the input, according to a timestamp *t* following a uniform distribution between 0 and 1. Below we describe the interpolation method for the amino acid type, the *C*_*α*_ coordinates, and the backbone orientations, respectively. The amino acid types at position *i* are denoted as ***s***_*i*_ and ***T*** _***s****i*_ for the ground truth CDR and the initial CDR, respectively, which are encoded as one-hot vectors. Therefore, calculating the interpolation simply requires computing the linear interpolation between the two vectors:

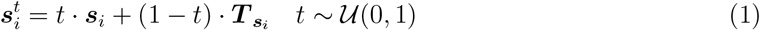

Similarly, the *C*_*α*_ atom coordinates are also linearly interpolated as:

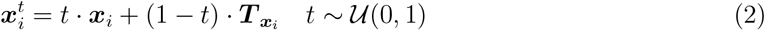

The interpolation of orientations differs slightly from the previous two cases. In order to ensure a smooth and uniform interpolation, we need to convert the orientations ***O***_***i***_ and ***T*** _***O****i*_ into quaternion representations: ***q***_*i*_ and 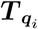. We then perform interpolation on the sphere using the spherical linear interpolation method, known as SLERP^26^:

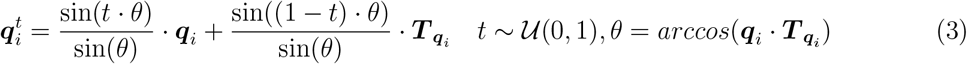

Therefore, for the *ℛ* = {***s***_*i*_, ***x***_*i*_, ***O***_*i*_| *i* = *l* + 1, …, *l* + *m*} in the dataset and a sampled *t*, following the method described above, we can obtain its interpolated representation, which is denoted as 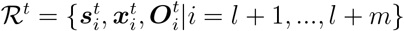

### Loss Function

The training loss comprises individual losses on the amino acid types, the *C*_*α*_ atom coordinates, and the orientations, which are calculated as follows. Regarding amino acid types, we directly employ the Mean Squared Error (MSE) on the vectorized representations:

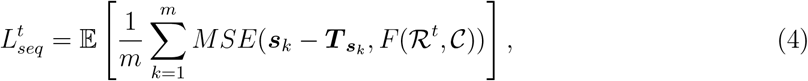

where *F* (*ℛ*^*t*^, *C*) denotes the predicted drift force on the vectorized representation of amino acid types. Correspondingly, for the *C*_*α*_ atom coordinates, due to their continuity, we also use the MSE for calculation:

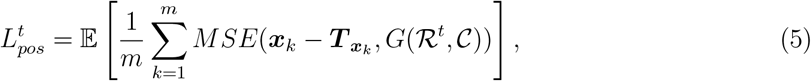

where *G*(*R*^*t*^,*C*) denotes the predicted drift force on the coordinates. The loss for orientations comprises two types of loss functions. First, we convert the orientation matrices into quaternions and implement Mean Squared Loss on them as follows:

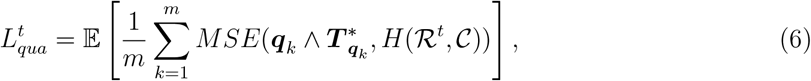

where the ∧ symbol indicates the Grassmann product between two quaternions, and the *symbol denotes the inverse of a quaternion. Then we exploit the loss function directly implemented on *SO*(3) the orientation matrices^11^:

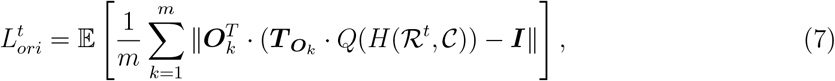

where *F* (*·*), *G*(*·*), and *H*(*·*) are neural networks responsible for predicting the drift force that maps the initial distribution to the target distribution. Additionally, *Q*(*·*) is the function mapping the quaternion representation to the *SO*(3) representation. By summing Eq.(4), (5), (6) and (7) and taking the expectation w.r.t *t*, we obtain the final training objective function:

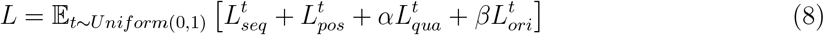

Here, both *α* and *β* range from 0 to 1, with their sum equaling one, balancing the two types of losses on the orientations.

### Parameterization with Neural Network

In this section, we briefly introduce the overall architecture of the network. The primary task of our network is to encode the input complex and predict the drift force that maps the initial distribution of amino acid types (*F*), *C*_*α*_ atom coordinates (*G*), and orientations (*H*) to the target distribution. The flow matching network incorporates a shared encoder for residue-level and pairwise representations, as well as three Multiple Layer Perceptron (MLP) heads for prediction of the drift forces on amino acid types, *C*_*α*_ atom coordinates, and orientations.

The residue-level input features include the amino acid type, all-atom coordinates, and side-chain torsional angles of each residue, which are processed into a vector *e*_*i*_ by a three-layer MLP with ReLU activation. The pairwise input features consist of the Euclidean distances and dihedral angles between each pair of residues, which are similarly encoded into vectors *z*_*ij*_ with another three-layer MLP. Both features are further concatenated with the embeddings of the flow matching time (t ∼ *U* (0, 1)) derived from two linear layers. Then, both *e*_*i*_ and *z*_*ij*_ are fed into an IPA module^27^, which is an orientation-aware and roto-translation invariant network, to obtain the contextual hidden representations *h*_*i*_ of each residue.

Next, three separate MLP heads operate on the hidden representations *h*_*i*_ to predict the drift forces for amino acid types, *C*_*α*_ atom coordinates, and orientations. For the probability distribution of amino acid types, a three-layer MLP with ReLU activation is utilized, which outputs 20-dimensional vectors for each residue, indicating the expected drift force of 20 canonical types of amino acids. For the *C*_*α*_ atom coordinates, a similar MLP is used to predict the three-dimensional scaled change in the local coordinate system, which is further projected to the global coordinate system with the interpolated orientation matrix:

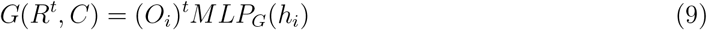

Such local-to-global transformations ensure the roto-translation equivariance of the predicted drift force on coordinates. For the residue orientations, we adopted the quaternion method^28^ which used four scalers to represent an orthogonal matrix. Therefore, an MLP with a four-dimensional output is utilized to predict the rotational drift force.

### Data Preparation

Our model is trained on SAbDab dataset^29^, which contains 13279 antibody-antigen complexes. We first clean up the data by discarding antibodies targeting non-protein antigens and structures with a resolution worse than 4Å. Next, to split the data for training, validation, and testing, we cluster all the antibodies in the dataset according to CDR-H3 sequences at 50% sequence identity using MMseqs2^24^. The test set comprises 20 non-redundant antibody-antigen complexes from different clusters. The complexes not in the same clusters as the test complexes are then partitioned into the training and the validation sets at a ratio of 20:1. In the training set, the average lengths of CDRs are as follows: CDR H1 is approximately 7.2, CDR H2 is 5.9, CDR H3 is 12.3, CDR L1 is 12.5, CDR L2 is 7.0, and CDR L3 is 9.2. In the test set, the average lengths are: CDR H1 is approximately 7.0, CDR H2 is 5.8, CDR H3 is 11.3, CDR L1 is 11.9, CDR L2 is 7.0, and CDR L3 is 9.2.

### Prior Distributions

Below we provide the implementations of the prior distributions tested in our experiments (see Figure 2). We followed the dataset splitting mentioned above for the distribution generated by the dyMEAN^8^ model. We fixed the sequences in the training data, excluding the antibody CDRs, and retrained the dyMEAN model. This model was then used to generate the initial CDRs for each complex in the dataset. For the random distribution, we directly leverage standard Gaussian for the vectorized representation of amino acid types. Regarding *C*_*α*_ atom coordinates, we rely on a standard Gaussian recentered at the centroid of the *C*_*α*_ atom coordinates from regions other than the CDR to be generated. As for the orientations, we initialize with ℐ 𝒢_*SO*(3)_, the isotropic Gaussian distribution on SO(3) parameterized by a mean rotation and a scalar variance30–^32^. In terms of the distribution generated by the kinematic-closure-based method KIC, we begin by discarding all structural information for the CDRs and using PyRosetta^33^ to sample *ϕ* and *ψ* angles of the residues within CDR on the Ramachandran plot^34^. Subsequently, we solve the kinematic equations to achieve closure of the entire structure and select the lowest energy structures to serve as our structural prior. During this process, we set our closure attempt to 50,000, established a minimum solution constraint of 1 for the kinematic equations, and filtered out all structures that experienced collisions. For each antibody, we generated 100 corresponding structures to serve as its prior distribution for training and inference. To generate the sequence prior distribution produced by ESM2, we mask all residues in the antibody CDRs, and then utilize the esm2_t30_150M_UR50D model to predict the sequences for the CDR regions.

### Model Training and Inference

In this section, we provide the implementation details for training and inference. During the training process, for the initialization of the distribution, we combine the dyMEAN distribution and the random distribution mentioned previously in Data Preparation by randomly selecting one method based on a probability *p*. Note that the probability *p* serves as a hyperparameter and can be adjusted to suit various requirements. To ensure the model does not get trapped in a local optimum and to guarantee the diversity of the initial distribution, we add a Gaussian noise when initializing the distribution with the dyMEAN model. Otherwise, the initial distribution would solidify into a few fixed samples, which would not be conducive to the generalization performance of the model. For *α* and *β* in the orientation loss, we set *α* to 1 and *β* to 0 for the initial 200,000 training iterations, and adjust *α* to 0 and *β* to 1 for the final 5,000 iterations. We provide further details in key resources table. We adopt Adam^35^ optimizer for model optimization. Algorithm 1 below presents the pseudocode of the overall training process of our model.

#### Algorithm 1

Training

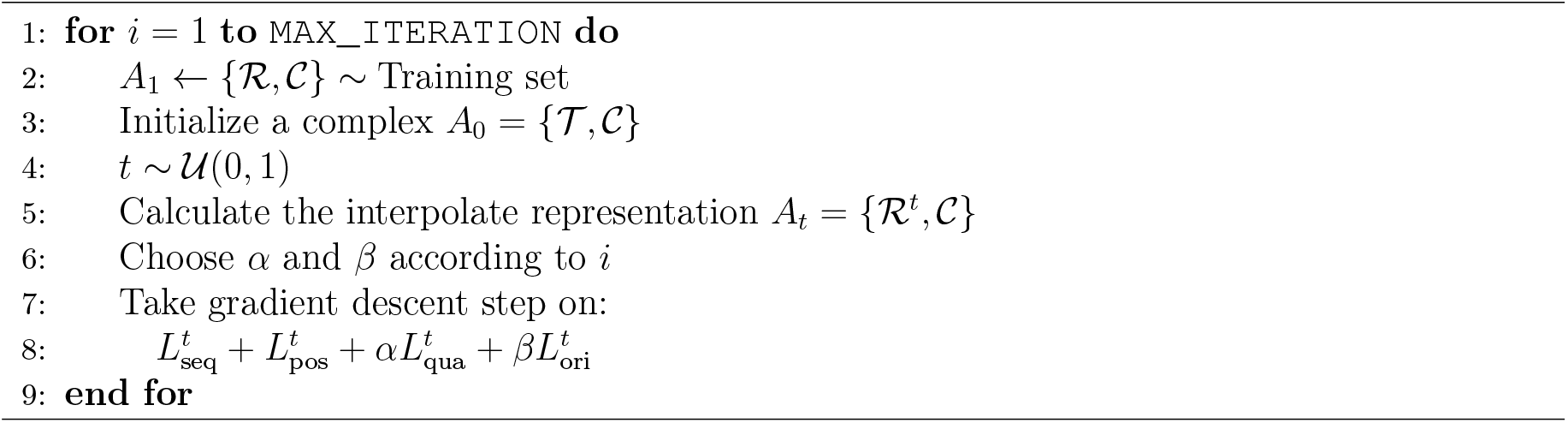

Next, we illustrate the details of the inference process. The training objective fundamentally aims at finding a transformation within a high-dimensional space that maps our initialized complex *A*_0_ directly to the real antibody *A*_1_. While it is feasible to achieve this transformation in a single step, such a direct transformation will likely lead to inferior performance in practical applications. Therefore, we employ the integral method, which decomposes the transformation into finer steps to enhance the accuracy. Specifically, we rescale the predicted drift force at each step by 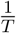and repeat the prediction-transformation cycle for *T* times. For the amino acid types and the *C*_*α*_ atom scoordinates, we can simply multiply the output of the neural network with 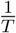 for transformations at each step.

For orientations, we explore two strategies during inference. The first strategy directly converts the quaternion predicted by our model into the rotational matrix, which is then applied to the input orientation matrix via right multiplication. The second strategy scales the angle of the rotation *θ* by 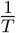 before the conversion of the quaternion. Our experiments demonstrate that the former method yields better results. Algorithm 2 below presents the pseudocode of the overall inference process of our model.

#### Algorithm 2

Inference

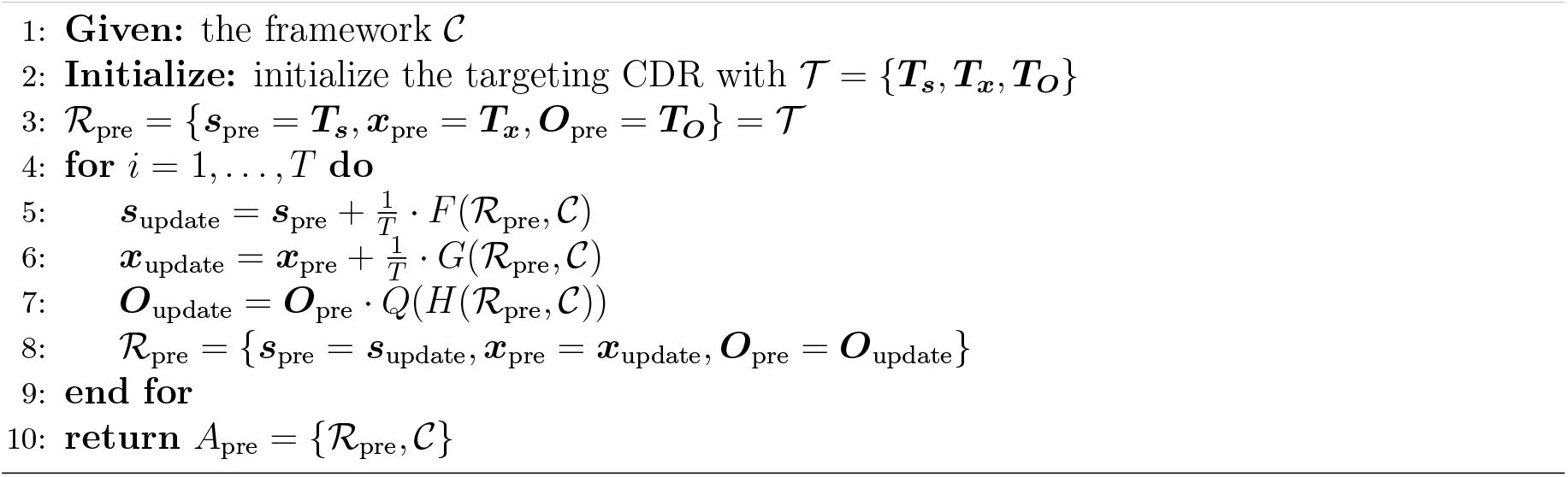

### Experimental procedure

In order to assess the potential of FlowDesign in generating high-affinity antibodies via cycles of yeast culture and fluorescence-activated cell sorting (FACS), our objective was to evolve the antibody Ibalizumab. The designed Ibalizumab HCDR3 library was codon-optimized for expression in yeast and cloned into pCTCON2 for galactose induction and surface expression in S.cerevisiae susing the Aga1p-Aga2p protein display system. The plasmid library was then transformed into the EBY100 strain using the Frozen Yeast Transformation Kit (Coolaber, SK2400). The constructed library was further cultured overnight in SD-CAA medium, followed by induction in SG-CAA medium. After induction, yeast cells were incubated with lab-made antigen CD4 d1-d2 for 1 hour and subsequently washed with PBS buffer. After incubation with fluorescently labeled antibodies, three consecutive rounds of sorting were performed on a BD Aria cell sorter. For the first round of FACS, 5 × 10_7_ induced yeast cells were centrifuged and washed with PBS buffer. And yeast cells were then incubated with 1 *µ*M biotin-conjugated CD4 d1-d4. After washing off unbound antigens, cells were then stained with 1:200 diluted anti-HA AF488 (Cell Signaling Technology cat. no. 2350) and Streptavidin APC (Miltenyi Biotec cat. no. 130-106-791) antibody for 1 hour at 4 °C. These cells were then centrifuged, resuspended in 3 ml of PBS buffer, and sorted. Sorting gates were defined by utilizing control cells displaying an antibody and stained with secondary antibodies, albeit without prior antigen incubation. The collection window is designed to isolate cells that secreted and displayed a functional antibody. In total, around 23,000 cells from 15,000,000 were collected and expanded. The second round of FACS was performed with similar conditions to the first; however, the antigen was changed to lab-made CD4 d1-d2, and the concentration of CD4 was reduced to 500 nM. For the second round, 3,000,000 cells were collected. For the last round of FACS, concentration of antigen was decreased to 200nM, while other conditions remained the same. In total, 3,000,000 cells were collected and plated on SD-CAA medium to isolate single clones. Forty-eight colonies were picked, cultured, induced and finally verified for binding with the antigen CD4 d1-d2. After three rounds of sorting, yeast cells were expanded to isolate yeast plasmids by using Yeast Plasmid Miniprep Kit (Zoman Biotechnology, ZP107). Yeast plasmids were then used as templates to amplify HCDR3 fragments to perform next-generation sequencing. After NGS analysis, the top 10 mutants were selected and expressed for characterization.

Antibodies targeting CD4 d1-d2 were expressed by cloning into pcDNA3.4 (biointron). For each antibody, 20mL of Expi293F cells (Thermo Fisher, A14527) were transfected with 20ug of plasmids. After one day, cells were enhanced with ExpiFectamine 293 Transfection Enhancer 1 and Enhance 2(Thermo Fisher, A14525). Cell supernatants were collected four days after transfection and each antibody was purified with 1mL rProtein A Beads(smart-life sciences, SA012100), which was then washed with 20 ml of PBS, eluted with 0.1M glycine (pH 3) and then neutralized with concentrated Tris buffer (pH 8). All antibodies were then dialyzed twice with PBS buffer.

Biolayer interferometry experiments for antibody binding assays were performed with an Octet R8 system (Sartorius). For antibody assays, Ibalizumab and its mutants were loaded onto protein A sensors (ForteBio, Pall LLC) at 10 *µ*g/ml for 120 s. The baseline interference was then read for 60 s in KB buffer (PBS, 0.1% BSA, 0.02%Tween). After the baseline measurement was obtained, antibodies were associated at 31.3 nM for 90 s, followed by an association KB buffer for 120s.

Pseudoviruses bearing the 11 HIV-1 envelopes (TRO_11, 398_F1_F6_20, CNE8, X2278_C2_B6, BJOX002000_03_2, 25710_2_43, CNE55, CNE4, CNE14, SF162 and JRFL) were generated by co-transfecting HEK293T cells with Env expression vectors and the pNL4–3R-Eluciferase viral backbone plasmid as described previously^36^. Pseudovirus-containing supernatants were collected 48 or 72 hours post transfection and the viral titers were measured by luciferase activity in relative light units (RLU) (Bright-Lite Luciferase Assay System, Vazyme, China). The supernatants were aliquoted and stored at -80°C until further use. Neutralization assays were performed by adding approximately 1.5×10_4_ GhostX4/R5 cells into 10 serial 1:3 dilutions of purified antibody starting from 7.5 *µ*g/ml. The mixture was then dispensed into a 96-well plate in duplicate and incubated for 1 hour at 37°C. The pseudovirus supernatants were then added and the cultures were maintained at 37°C for an additional 48h before luciferase activity was measured. Neutralizing activity was measured by the reduction in luciferase activity compared to the controls. The half-maximal inhibitory concentrations (IC50), the concentrations required to inhibit infection by 50% compared to the controls, were calculated using the dose-response-inhibition model with a 5-parameter Hill slope equation in GraphPad Prism 10(GraphPad Software, USA.).

## SUPPLEMENTAL INFORMATION

**Figure S1.**
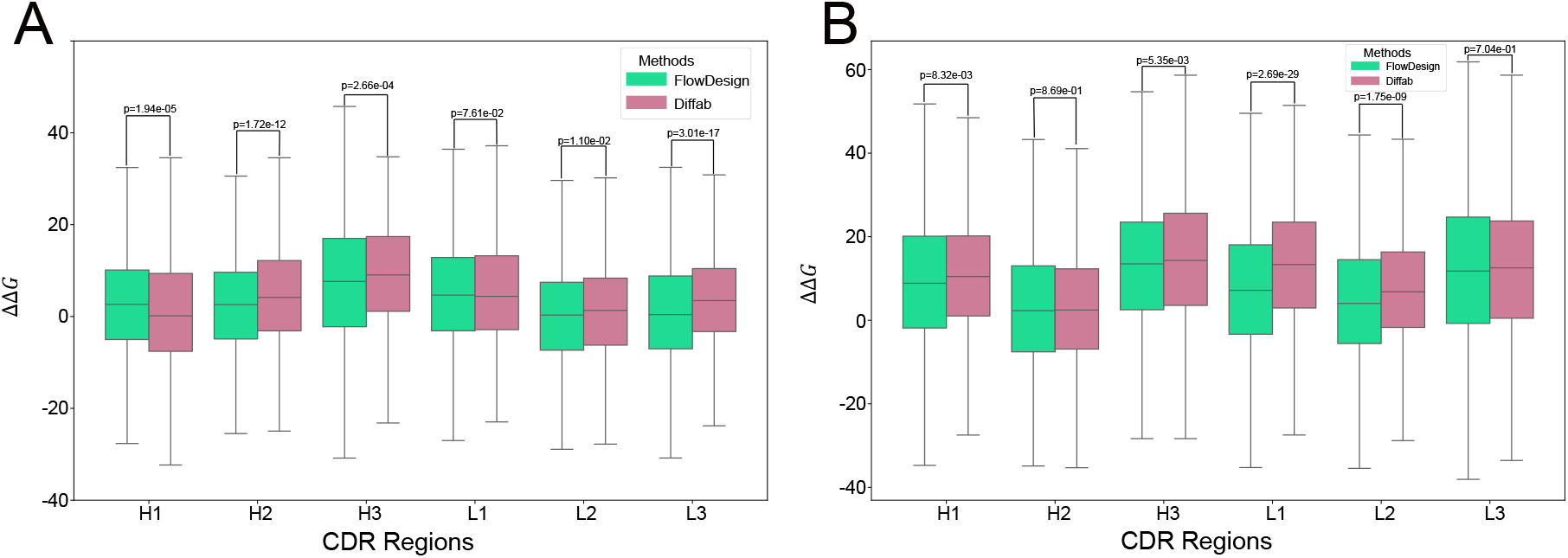
Comparison of ΔΔ*G* for different side chain generation strategies and energy calculation methods on different CDRs. **(A)** ΔΔ*G* comparison using OpenMM^37^ for side-chain packing. **(B)** ΔΔ*G* comparsion using FoldX^38^ for binding energy calculation.

**Figure S2.**
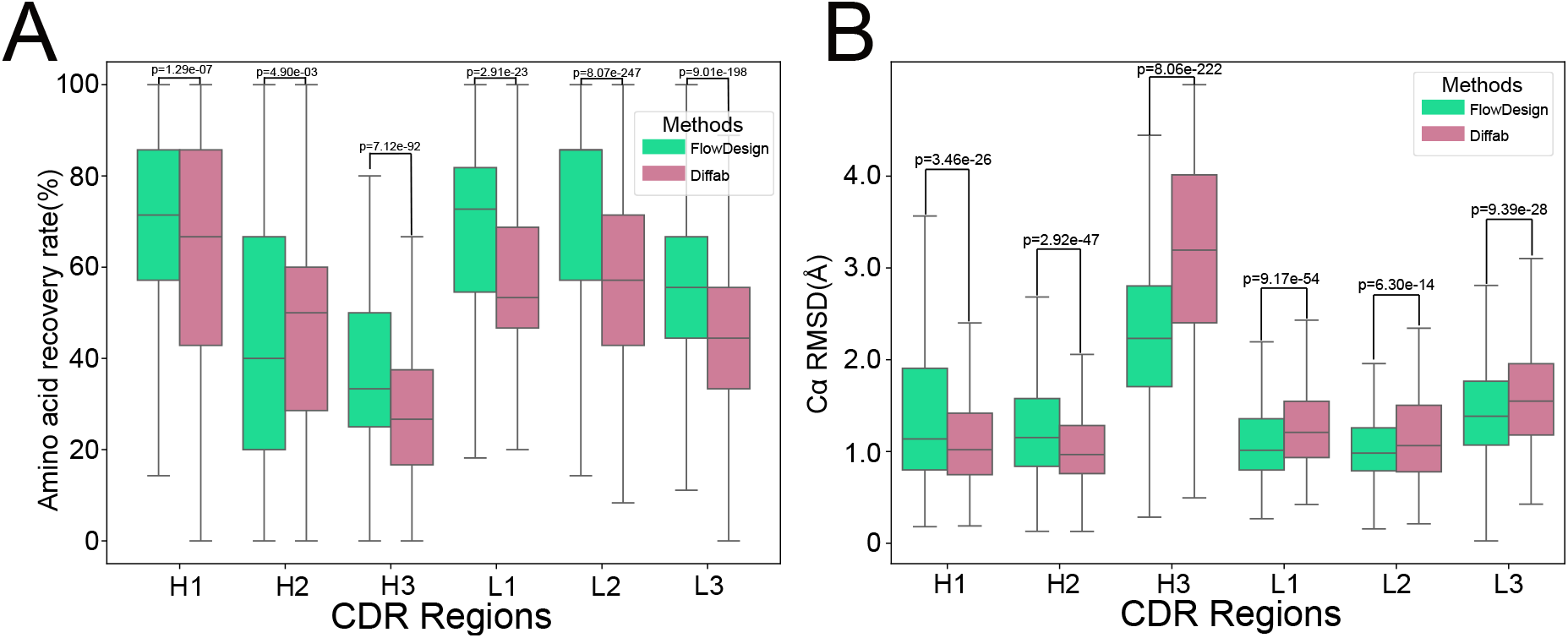
Comparison of performance between randomly initialized FlowDesign and Diffab on different CDRs. **(A)** Comparison of the distribution of AAR in generated conformations. **(B)** Comparison of the distribution of RMSD in generated conformations.

**Figure S3.**
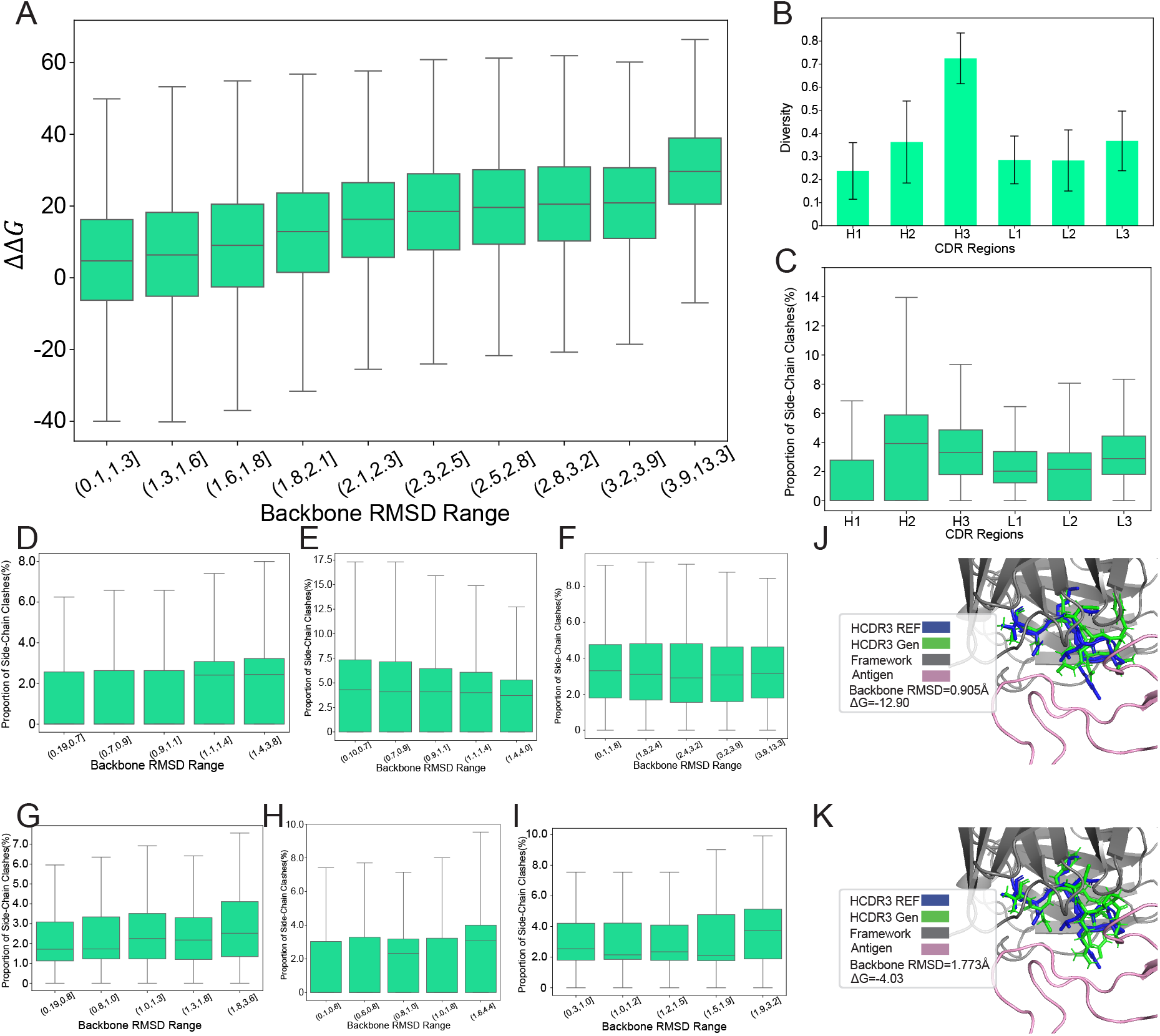
**(A)** The relationship between the RMSD of the generated conformations’ backbone and energy. **(B)** The diversity of sequences in the generated conformations. **(C)** Proportion of side-chain clashes in different CDRs. **(D)-(I)** The relationship between the RMSD of the generated conformations’ backbone and side-chain clashes in different CDRs. In order: CDR H1, H2, H3, L1, L2, L3. **(J)(K)** Visualization of generated conformations and their side-chains.

**Figure S4.**
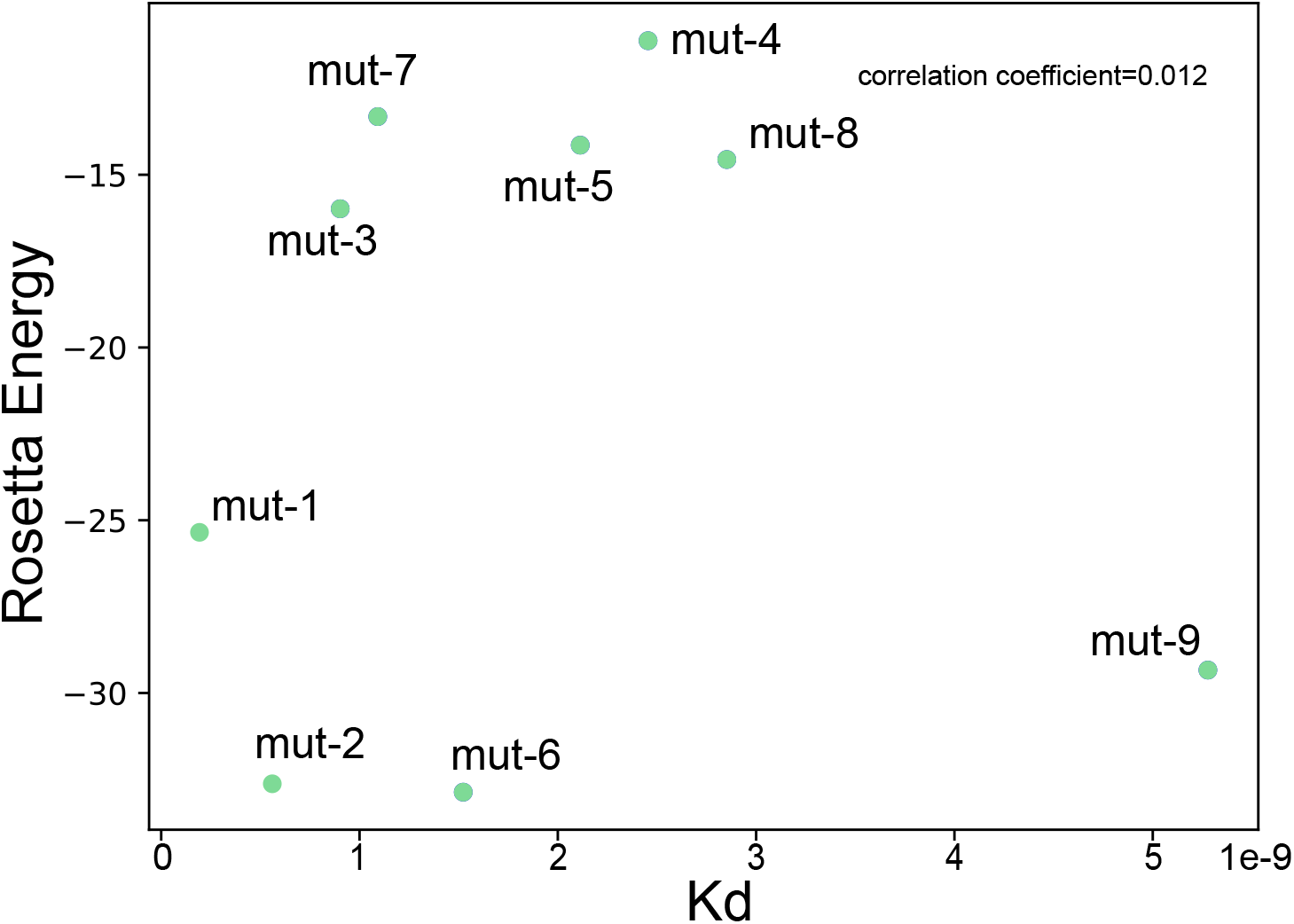
The relationship between the Kd of HIV mutant antibodies and their corresponding Rosetta energy.

**Figure S5.**
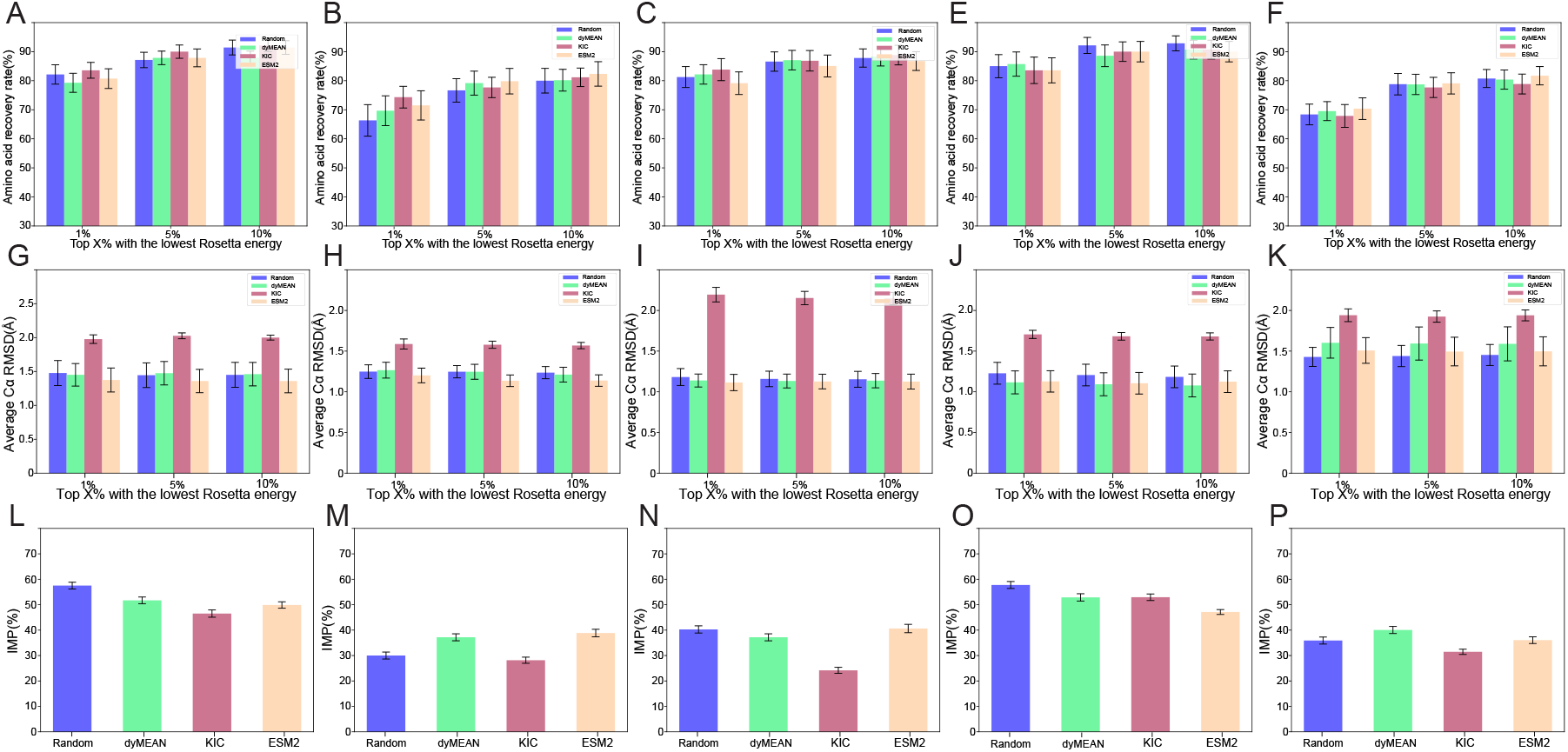
**(A)-(F)** Performance for amino acid recovery rates using different prior distributions in CDR H1, H2, L1, L2, and L3. **(G)-(K)** Performance for average *C*_*α*_ RMSD using different prior distributions in CDR H1, H2, L1, L2, and L3. **(L)-(P)** Performance for IMP using different prior distributions in CDR H1, H2, L1, L2, and L3.

**Figure S6.**
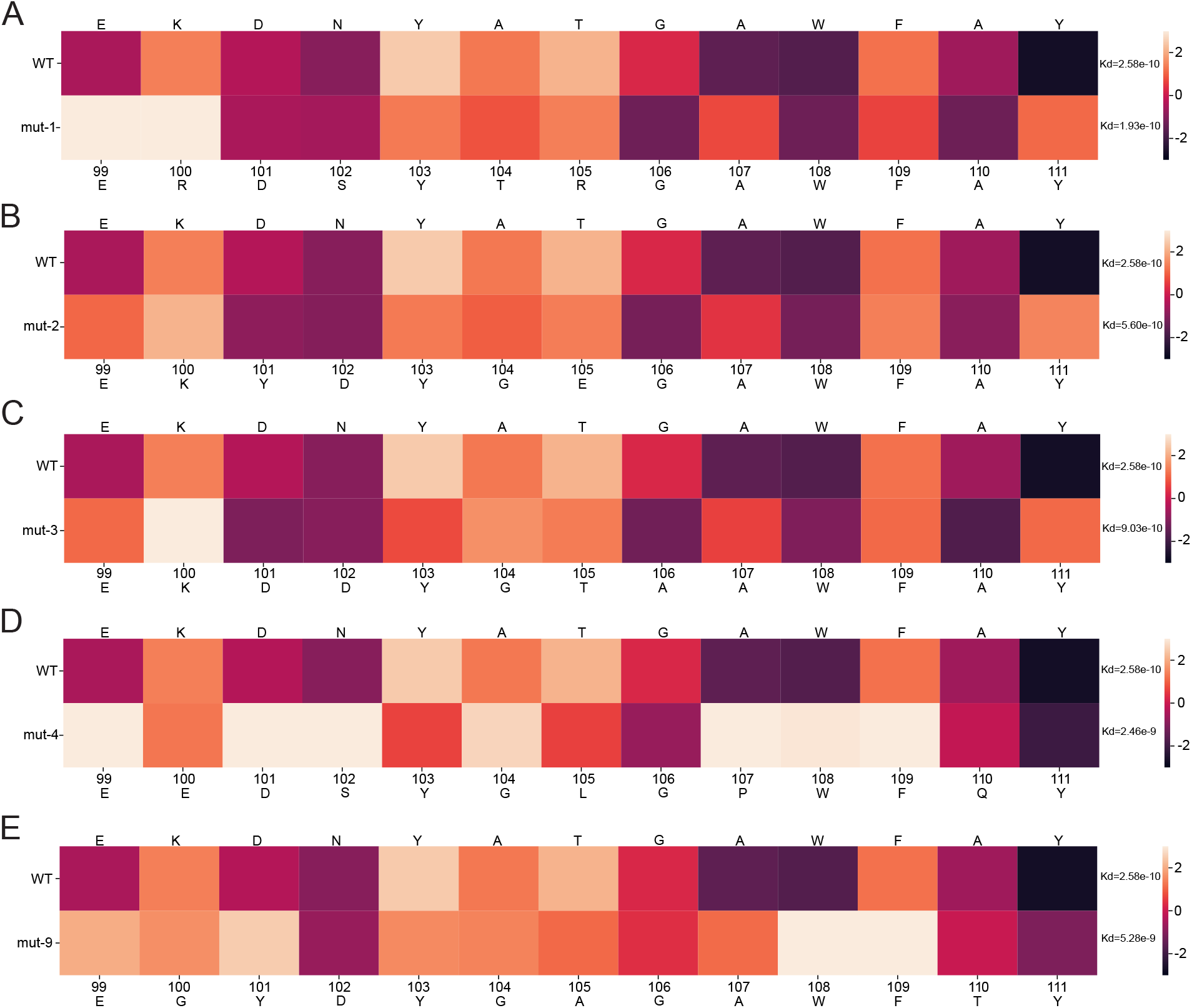
**(A)-(E)** Visualizations of the energy change for each residue in the CDRH3 of imab-mut 1, 2, 3, 4, and 9 relative to imab-WT, along with their corresponding Kd values.

